## Supporting File 1 for "Activation of cryptic splicing in bovine *WDR19* is associated with reduced semen quality and male fertility"

**Supporting Table 1: Phenotypic correlations (off-diagonal) and heritability (diagonal) of the traits studied**

|  | VOL | CONC | MOTIL | ANOHEAD | ANOTAIL | NTARGET | FERT |
| --- | --- | --- | --- | --- | --- | --- | --- |
| VOL | 0.27 ± 0.05 | -0.37 ± 0.03 | -0.17 ± 0.04 | 0.11 ± 0.04 | 0.12 ± 0.04 | 0.15 ± 0.04 | -0.13 ± 0.04 |
| CONC |  | 0.25 ± 0.06 | 0.06 ± 0.04 | -0.02 ± 0.04 | -0.02 ± 0.04 | 0.01 ± 0.04 | 0.00 ± 0.04 |
| MOTIL |  |  | 0.10 ± 0.04 | -0.67 ± 0.03 | -0.75 ± 0.02 | -0.61 ± 0.03 | 0.22 ± 0.04 |
| ANOHEAD |  |  |  | 0 | 0.84 ± 0.02 | 0.58 ± 0.03 | -0.21 ± 0.04 |
| ANOTAIL |  |  |  |  | 0.02 ± 0.03 | 0.68 ± 0.03 | -0.21 ± 0.04 |
| NTARGET |  |  |  |  |  | 0.12 ± 0.06 | -0.20 ± 0.04 |
| FERT |  |  |  |  |  |  | NA |

VOL – ejaculate volume; CONC – sperm concentration; MOTIL – sperm motility; ANOHEAD – proportion of sperm with head anomalies; ANOTAIL – proportion of sperm with tail anomalies; NTARGET – number of sperm per straw; FERT – bull fertility
