## Supplementary figures and images for "Activation of cryptic splicing in bovine *WDR19* is associated with reduced semen quality and male fertility"

### Supporting File 2

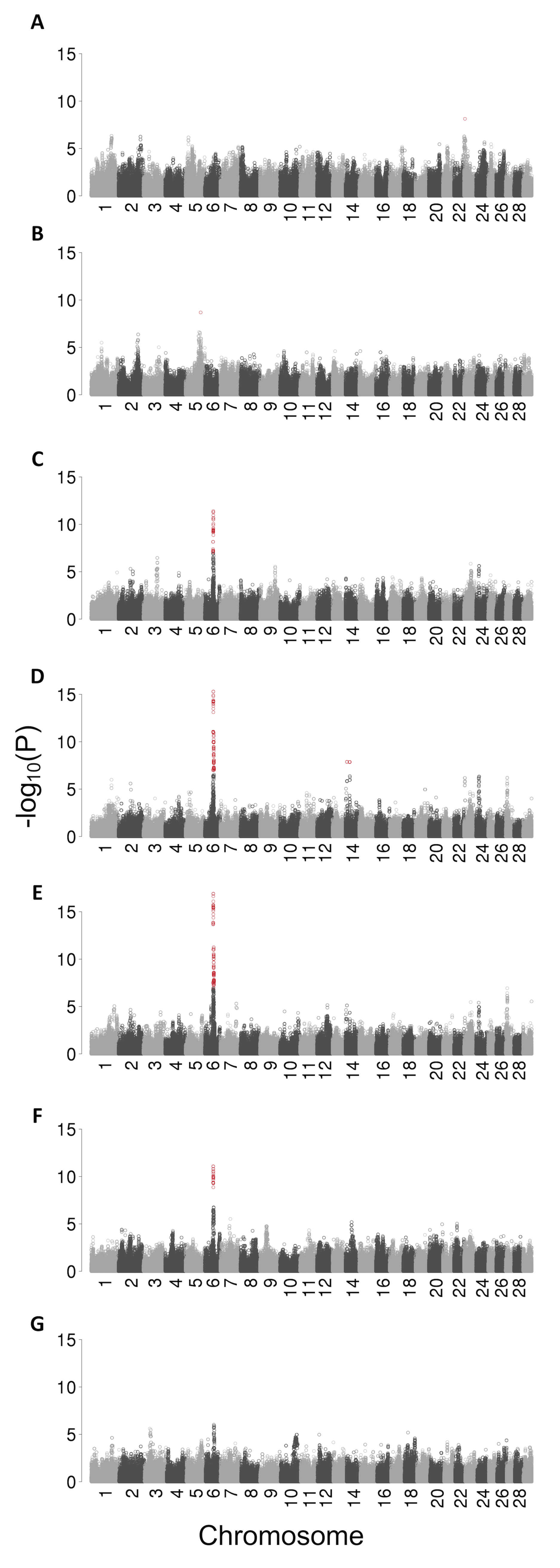

### Supporting File 3

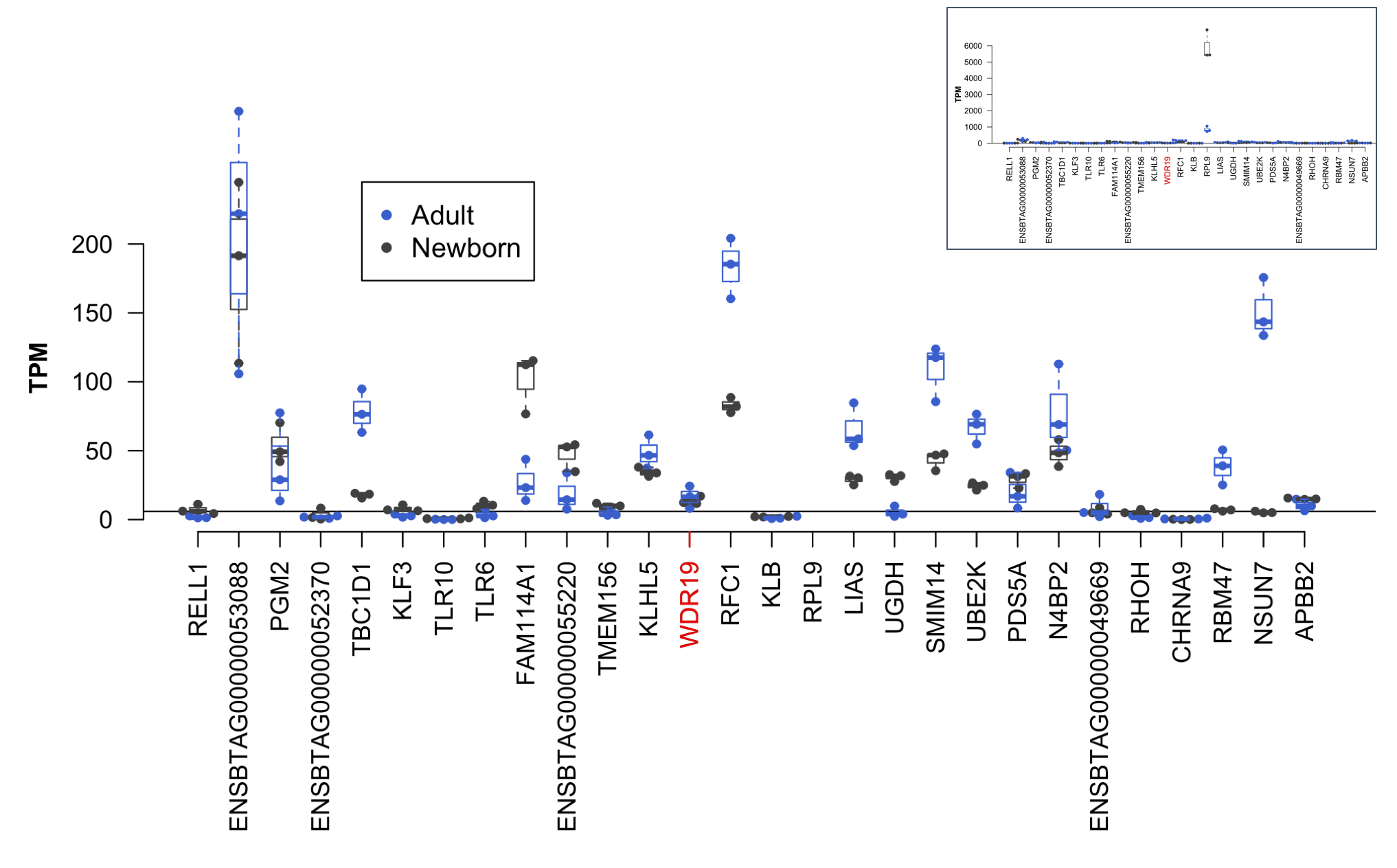

### Supporting File 7

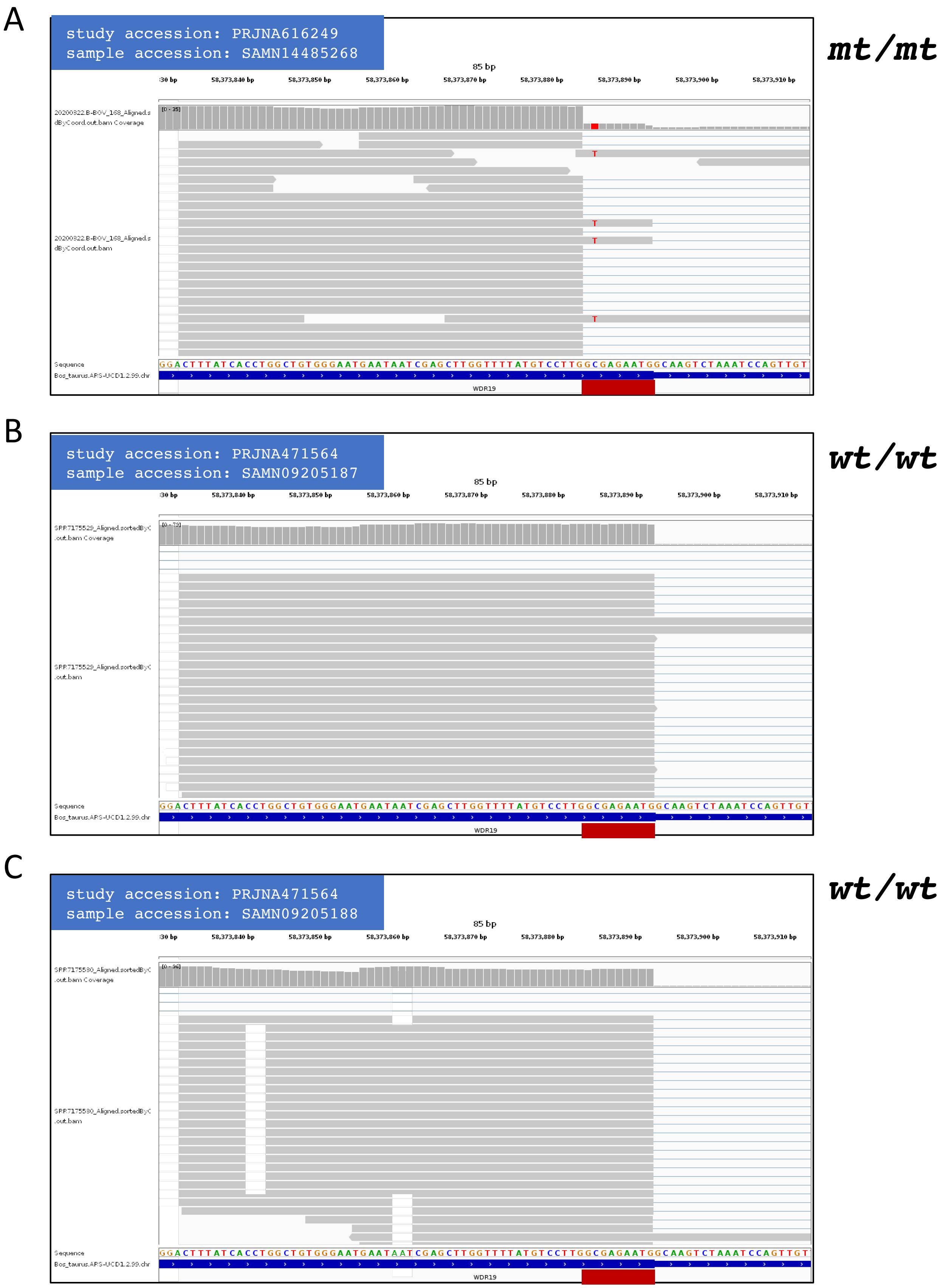

### Supporting File 8

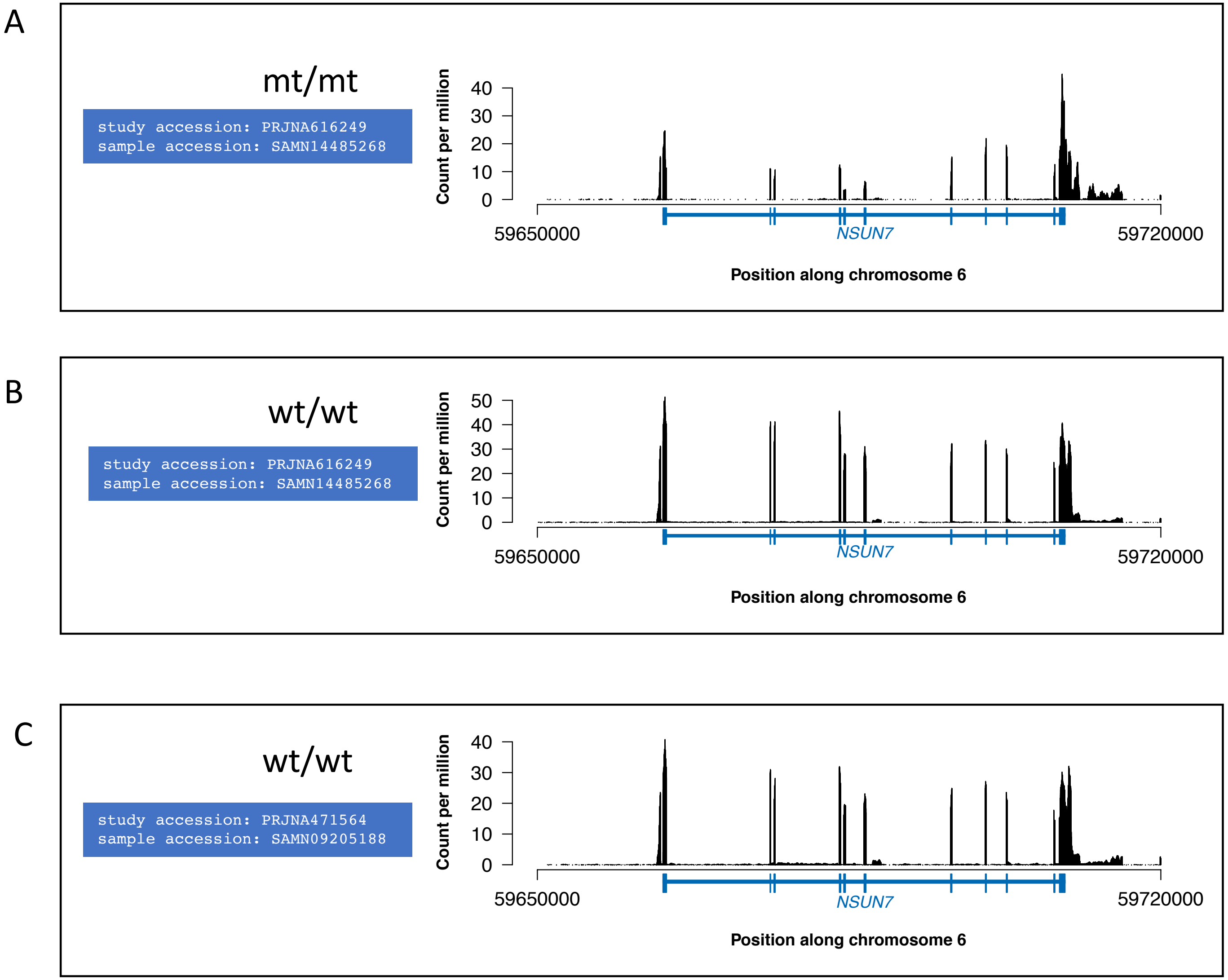
