## Supporting File 10 for "Activation of cryptic splicing in bovine *WDR19* is associated with reduced semen quality and male fertility": GWAS.html

Untitled2


### R syntax to plot the results of the haplotype-based GWAS for sperm motility¶

In [1]:

```
# read the data, assuming 
#   a) the R script haplo_linear.R ran successfully 
#   b) the output file GWAS_motil_6 is located in $HOME

home_directory <- system(paste("echo $HOME"), intern=TRUE)

file <- read.table(paste(home_directory, "/GWAS_motil_6", sep=""),
                   header=TRUE, colClasses=c(rep("numeric", 5), "character", rep("numeric", 7), "character")
                  )
head(file)
```

| CHR | startSNP | stopSNP | startPosition | stopPosition | Haplotype | fq | AA | AB | BB | beta | se\_beta | pval | test |
| --- | --- | --- | --- | --- | --- | --- | --- | --- | --- | --- | --- | --- | --- |
| 6 | 1 | 51 | 123308 | 386837 | 010011110001100010111000010111101111001111111110011 | 0.09697733 | 647 | 140 | 7 | 0.26523384 | 0.2023334 | 0.1902838 | ADD |
| 6 | 1 | 51 | 123308 | 386837 | 011111111111110111101000011110100101001111111110011 | 0.03841310 | 735 | 57 | 2 | -0.14959426 | 0.3198791 | 0.6401586 | ADD |
| 6 | 1 | 51 | 123308 | 386837 | 101110000000111111111111111111111111111111111111111 | 0.13853904 | 584 | 200 | 10 | 0.16909176 | 0.1811798 | 0.3509619 | ADD |
| 6 | 1 | 51 | 123308 | 386837 | 110011111111010111110001011110100001000000001000000 | 0.03400504 | 742 | 50 | 2 | -0.21465488 | 0.3305047 | 0.5162210 | ADD |
| 6 | 1 | 51 | 123308 | 386837 | 111111111111110111110000010111101111001111111110011 | 0.09571788 | 652 | 132 | 10 | 0.04393742 | 0.1972449 | 0.8237837 | ADD |
| 6 | 1 | 51 | 123308 | 386837 | 111111111111111111111101011111101111011010001110011 | 0.01007557 | 778 | 16 | 0 | 0.75740438 | 0.5868904 | 0.1972444 | ADD |

In [2]:

```
# extract the statistics for the recessive tests
file_rec <- file[file$test=="REC", ]
```

In [3]:

```
# inspect the positions of the most significantly associated haplotypes
head(file_rec[order(file_rec$pval, decreasing=FALSE), ])[, -c(2, 3, ncol(file_rec))]

# the top haplotype occurs in the homozygous state in 46 bulls (column BB)
# its frequency in 794 bulls is 0.2411839 (column BB)
# the top haplotypes extend from 57538068 to 57538068 bp and their P-value is 2.540683e-27
```

|  | CHR | startPosition | stopPosition | Haplotype | fq | AA | AB | BB | beta | se\_beta | pval |
| --- | --- | --- | --- | --- | --- | --- | --- | --- | --- | --- | --- |
| 28557 | 6 | 57538068 | 57939675 | 111111000111100000000000111111111111111111110000000 | 0.2411839 | 457 | 291 | 46 | -3.712953 | 0.3300589 | 2.540683e-27 |
| 28559 | 6 | 57614247 | 57993128 | 000000000111111111111111111110000000000011111111111 | 0.2411839 | 457 | 291 | 46 | -3.712953 | 0.3300589 | 2.540683e-27 |
| 28552 | 6 | 57335668 | 57624763 | 111111111111111111111111110111111111000111100000000 | 0.2386650 | 460 | 289 | 45 | -3.673123 | 0.3352339 | 4.298359e-26 |
| 28569 | 6 | 57985954 | 58096920 | 111111101111010000111111111111111111111111111111101 | 0.2518892 | 444 | 300 | 50 | -3.355041 | 0.3200249 | 3.724616e-24 |
| 28572 | 6 | 58028433 | 58115812 | 000111111111111111111111111111111101111000000000111 | 0.2518892 | 444 | 300 | 50 | -3.355041 | 0.3200249 | 3.724616e-24 |
| 28578 | 6 | 58085235 | 58217062 | 111101111000000000111111111111111111111111111111111 | 0.2506297 | 446 | 298 | 50 | -3.355041 | 0.3200249 | 3.724616e-24 |

In [4]:

```
# plot the results of the association study

plot((file_rec$startPosition + file_rec$stopPosition)/2/1000000, -log10(file_rec$pval),
    xlab="Position (in Mb)", ylab="-log10(P)", pch=23, col="grey25", bg="#0069B4", lwd=.3,
    las=1)
```
