## Supporting File 10 for "Activation of cryptic splicing in bovine *WDR19* is associated with reduced semen quality and male fertility": haplo_linear.R

```
#+++++++++++++++++++++++++++++++++++++++++++++++++++++++#	
#														#
#	Association testing between haplotypes and			#
#	quantitative traits									#
#														#
#	Hubert Pausch			#  
#	06.12.2019											#
#														#
#														#
#  This R script requires following files:				#
#    - phenotype file in plink format					#
#    - haplotype file in MaCH/Minimac format			#
#    - map-file in plink format							#
#    - eigenvector file in GCTA format					#
#														#
#														#	
#+++++++++++++++++++++++++++++++++++++++++++++++++++++++#


args <- commandArgs(TRUE)

chr <- args[1]
phenotype_file <- args[2]
haplotype_file <- args[3]
mapposition_file <- args[4]
eigenvec_file <- args[5]
number_evecs <- as.numeric(args[6])
sliding_window <- as.numeric(args[7])
haplo_length <- as.numeric(args[8])
min_haplo_freq <- as.numeric(args[9])
outputfile <- args[10]

# minimum proportion of homozygous animals for recessive tests:
min_freq_hom <- 0.01

cat(paste("\n reading data ... \n"))

pheno <- read.table(phenotype_file, colClasses=c(rep('character', 2), 'numeric'))
haplo_file <- read.table(haplotype_file, colClasses='character')
map_file <- read.table(mapposition_file, colClasses=c('numeric','character', rep('numeric', 2)))
evec_file <- read.table(eigenvec_file, colClasses='character')

for (i in 3:ncol(evec_file)){
	evec_file[,i] <- as.numeric(evec_file[,i])
}

evec_file <- evec_file[, 1:(number_evecs+2)]

names(pheno)[2] <- names(haplo_file)[1] <- names(evec_file)[2] <- 'ID'
pheno <- pheno[!is.na(pheno[,3]),]; pheno <- pheno[pheno[,3]!='9999',]


haplo_file[,1] <- unlist(strsplit(haplo_file[ ,1], '->'))[seq(from=2, by=2, to=nrow(haplo_file)*2)]
haplo_file <- merge(haplo_file, pheno, by='ID')
haplo_file <- merge(haplo_file, evec_file[ ,c(2:number_evecs)], by='ID')
evecs <- haplo_file[seq(from=1, by=2, to=nrow(haplo_file)), c(6:ncol(haplo_file))]
haplo_file <- haplo_file[ ,c(1:5)]
phenotype <- haplo_file[seq(from=1, by=2, to=nrow(haplo_file)), 5]

min_haplos <- round(min_haplo_freq * nrow(haplo_file), 0)

ht1 <- seq(from=1, by=2, to=nrow(haplo_file))
ht2 <- seq(from=2, by=2, to=nrow(haplo_file))

n_snps <- nchar(haplo_file[1, 3])
window_starts <- sort(unique(c(seq(from=1, by=sliding_window, to=(n_snps-haplo_length+sliding_window)), n_snps-haplo_length)))

collect_results_add <- numeric(0)
collect_results_rec <- numeric(0)
collect_results_dom <- numeric(0)

cat(paste("\n start the association testing for ", length(window_starts[window_starts+haplo_length <= n_snps]), " haplotype windows on chromsome ", chr, "\n\n", sep=""))

for (i in 1:length(window_starts[window_starts+haplo_length <= n_snps])){
	
	if(i %% 100 == 0) {
		cat(paste("\r haplotype windows processed:   ", i,  sep=""))
	}
	
	hts <- substr(haplo_file[,3], window_starts[i], window_starts[i]+haplo_length)

	if (length(which(table(hts)>=min_haplos)) >= 1){
	
		for (j in 1:length(names(which(table(hts) >= min_haplos)))){

			#additive tests
			test_ht <- names(which(table(hts) >= min_haplos))[j]
			fq <- length(hts[hts==test_ht])/length(hts)
			count <- rep(0, length(ht1))
			count[hts[ht1]==test_ht] <- 1
			count[hts[ht2]==test_ht] <- count[hts[ht2]==test_ht]+1
			model <- summary(lm(phenotype~count+as.matrix(evecs)))
			stats_ht <- c(i,j, test_ht, fq, length(count[count==0]),length(count[count==1]),length(count[count==2]), model$coefficients[2,1], model$coefficients[2,2], model$coefficients[2,4])
			collect_results_add <- rbind(collect_results_add, stats_ht)
			
			# recessive tests
            if (length(count[count==2])>=1){
            	if ((length(count[count==2]) / length(count)) > min_freq_hom){
					rec <- rep(0, length(count))
					rec[count==2] <- 1
					model <- summary(lm(phenotype~rec+as.matrix(evecs)))
					stats_ht <- c(i,j, test_ht, fq, length(count[count==0]),length(count[count==1]),length(count[count==2]), model$coefficients[2,1], model$coefficients[2,2], model$coefficients[2,4])
					collect_results_rec <- rbind(collect_results_rec, stats_ht)
        	    }
			}
			
			# dominant tests
			if (length(count[count==2])>=1){
				dom <- rep(1, length(count))
				dom[count==0] <- 0
				model <- summary(lm(phenotype~dom+as.matrix(evecs)))
				stats_ht <- c(i,j, test_ht, fq, length(count[count==0]),length(count[count==1]),length(count[count==2]), model$coefficients[2,1], model$coefficients[2,2], model$coefficients[2,4])
				collect_results_dom <- rbind(collect_results_dom, stats_ht)
    
            }
        }
    }
}

cat(paste("\r haplotype windows processed:   ", i,  sep=""))


outfile_add <- as.data.frame(cbind(rep(chr,nrow(collect_results_add)), window_starts[as.numeric(collect_results_add[,1])], window_starts[as.numeric(collect_results_add[,1])]+haplo_length, map_file[window_starts[as.numeric(collect_results_add[,1])],4], map_file[window_starts[as.numeric(collect_results_add[,1])]+haplo_length,4], collect_results_add[,3], collect_results_add[,4] , collect_results_add[,5], collect_results_add[,6], collect_results_add[,7], collect_results_add[,8], collect_results_add[,9], collect_results_add[,10], c("ADD")), row.names=c(1:nrow(collect_results_add)), stringsAsFactors=FALSE)
outfile_dom <- as.data.frame(cbind(rep(chr,nrow(collect_results_dom)), window_starts[as.numeric(collect_results_dom[,1])], window_starts[as.numeric(collect_results_dom[,1])]+haplo_length, map_file[window_starts[as.numeric(collect_results_dom[,1])],4], map_file[window_starts[as.numeric(collect_results_dom[,1])]+haplo_length,4], collect_results_dom[,3], collect_results_dom[,4] , collect_results_dom[,5], collect_results_dom[,6], collect_results_dom[,7], collect_results_dom[,8], collect_results_dom[,9], collect_results_dom[,10], c("DOM")), row.names=c(1:nrow(collect_results_dom)), stringsAsFactors=FALSE)
outfile_rec <- as.data.frame(cbind(rep(chr,nrow(collect_results_rec)), window_starts[as.numeric(collect_results_rec[,1])], window_starts[as.numeric(collect_results_rec[,1])]+haplo_length, map_file[window_starts[as.numeric(collect_results_rec[,1])],4], map_file[window_starts[as.numeric(collect_results_rec[,1])]+haplo_length,4], collect_results_rec[,3], collect_results_rec[,4] , collect_results_rec[,5], collect_results_rec[,6], collect_results_rec[,7], collect_results_rec[,8], collect_results_rec[,9], collect_results_rec[,10], c("REC")), row.names=c(1:nrow(collect_results_rec)), stringsAsFactors=FALSE)
outfile <- rbind(outfile_add, outfile_dom, outfile_rec)

colnames(outfile) <- c('CHR', 'startSNP', 'stopSNP', 'startPosition', 'stopPosition', 'Haplotype', 'fq', 'AA', 'AB', 'BB', 'beta', 'se_beta', 'pval', 'test')

write.table(outfile, paste(outputfile, "_", chr, sep=""), row.names=FALSE, col.names=TRUE, quote=FALSE, sep="\t")


cat(paste("\n\n++++++++++\n output written to: ", paste(outputfile, "_", chr, sep=""), "\n++++++++++\n Program finished\n\n"))
```
