## Supporting File 10 for "Activation of cryptic splicing in bovine *WDR19* is associated with reduced semen quality and male fertility": README

README

#	In order to carry out the GWAS, please adjust the paths accordingly:
#	+++++++++++++++++++++++++++++++++++++
path_to_Rscript=$HOME/Supporting_File_10
path_to_data=$HOME/Supporting_File_9
path_to_output=$HOME
#	+++++++++++++++++++++++++++++++++++++


# the following variables specify the input files
#	+++++++++++++++++++++++++++++++++++++
chr=6
input_file_haplotype=${path_to_data}/haplotypes_${chr}
input_file_map=${path_to_data}/markers_${chr}
input_file_phenotype=${path_to_data}/motil
input_file_evecs=${path_to_data}/evecs
#	+++++++++++++++++++++++++++++++++++++

# the following variables specify model parameters
#	+++++++++++++++++++++++++++++++++++++
n_evecs=10
sliding_window=15
haplo_length=50
min_haplo_freq=0.01
#	+++++++++++++++++++++++++++++++++++++

#	+++++++++++++++++++++++++++++++++++++
output_prefix=${path_to_output}/GWAS_motil


# the following command envokes the haplotype-based GWAS in a unix environment, assuming that R is installed and in the $PATH
#	+++++++++++++++++++++++++++++++++++++
Rscript ${path_to_Rscript}/haplo_linear.R ${chr} ${input_file_phenotype} ${input_file_haplotype} ${input_file_map} ${input_file_evecs} ${n_evecs} ${sliding_window} ${haplo_length} ${min_haplo_freq} ${output_prefix}
